# Reconstructing regulatory pathways by systematically mapping protein localization interdependency networks

**DOI:** 10.1101/116749

**Authors:** James Dodgson, Anatole Chessel, Federico Vaggi, Marco Giordan, Miki Yamamoto, Kunio Arai, Marisa Madrid, Marco Geymonat, Juan Francisco Abenza, José Cansado, Masamitsu Sato, Attila Csikasz-Nagy, Rafael Edgardo Carazo Salas

## Abstract

A key goal of functional genomics is to elucidate how genes and proteins act together in space and time, wired as pathways, to control specific aspects of cell biological function. Here, we develop a method to quantitatively determine proteins’ localization interdependencies at high throughput. We show that this method can be used to systematically obtain weighted, signed and directional pathway relationships, and hence to reconstruct a detailed pathway wiring. As proof-of-principle, we focus on 42 factors that control cell polarity in fission yeast *(Schizosaccharomyces pombe)* and use high-throughput confocal microscopy and quantitative image analysis to reconstruct their Localization Interdependency Network (LIN). Through this approach we identify 554 pairwise interactions across the factors, including 98% putative new directed links. Validation of an unexpected interaction between two polarity factor subgroups - the polarity landmark proteins and the cell integrity pathway components - by orthogonal phenotyping demonstrates the power of the LIN approach in detecting subtle, systems-level causal connections.

## INTRODUCTION

Cells are controlled not by genes acting in isolation but by genes and proteins that interact in complex regulatory networks. For the past two decades, network biology^1^ has combined high-throughput biology and computational approaches to build exhaustive and unbiased maps of those interaction networks, by interactome analysis^2,3^ or by genetics or epistasis approaches^4-7^. Those efforts however are based on ‘bulk’ readouts obtained from populations of cells and hence ignore the dynamical nature of gene and protein interactions, which take place at specific times and locations in specific cells. That level of information can only be accessed by singlecell approaches, looking at molecules in their native environment.

Here we developed a method to infer interaction networks by quantitatively exploiting proteins’ spatiotemporal localization pattern in live cells, using large-scale high-content microscopy^8-11^. The method consists in establishing whether the subcellular localization of a protein **A** is affected significantly by deregulation of another protein **B**, to obtain an edge **B ➔ A** (‘disruption of **B** influences the localization pattern of **A**’), and combining the significant, statistically-weighted and directed edges for all proteins in a group to obtain the proteins’ Localization Interdependency Network (LIN). This is done in practice by imaging a combinatorial library of cell lines in which the localization pattern of every protein - labelled with GFP - can be assessed quantitatively upon disruption of every other protein - for example, by knockout (KO) of their corresponding gene - across populations of cells, using automated image processing.

By building weighted, signed and directed networks based on complex quantitative readouts from single cells, the LIN method leverages the power of both network biology and single cell phenotyping, allowing refined identification of spatiotemporally-resolved causality between proteins in pathways.

## RESULTS

### Implementation of the LIN method

As example biological system to implement the LIN method, we choose the molecular pathway that controls polarized cellular growth in fission yeast (*Schizosaccharomyces pombe*). *S. pombe* cells have a simple cylindrical shape and a stereotypical growth pattern through the cell cycle that is controlled by well-studied genes and proteins (‘polarity factors’) ^12-14^ Though little information is available on how the polarity factors are functionally connected to one another as an overall network^12,13^ many of them localize to the growing areas of cells, which are typically restricted to the ends of the cylindrical cells, where they are enriched. This provides an opportunity to exploit their distinct localization pattern to infer network information.

Therefore and as a starting point, we selected a subset of 42 *S. pombe* polarity factors by bioinformatics analysis^15^ and literature mining, and generated 42 cell lines coexpressing each factor GFP-tagged together with the cell growth marker RFP-Bgs4 (Bgs4 is a (1,3)β-D-glucan cell wall synthase^16^; for this and other methodological details see Methods). Most of the GFP-tagged cell lines were specifically created for this study allowing relative uniformity of composition, and optimization of tagging to the N- or C-terminal domains. Where possible, we sought to generate cell lines where GFP-tagged factors were expressed under the control of their endogenous promoters (32/42 factors), to minimize over-expression artifacts; alternatively, the GFP-tagged factors were driven by an inducible nmt-promoter (Maundrell, 1990, Basi et al. 1993) (Supplementary table 1; note: in all GFP-tagged cell lines the only copy of the polarity factor was the GFP-tagged copy). In parallel, we compiled 29 cell lines with each non-essential polarity factor gene knocked out (most of them available from Bioneer, Korea^17^; note: 14 of the 42 factors selected are essential or sterile and KO cell lines for them could not be made).

We then crossed the 42 GFP cell lines against the 29 KO cell lines to obtain a combinatorial library of 1,232 haploid lines, where the localization of each GFP-labelled polarity factor at growing cell domains could be monitored in absence of every factor knocked out. The library was arrayed in 96-microwell plates, arranged so that each plate contained KO cell lines with a common GFP-labelled polarity factor (Supplementary Figure 1). This arrangement had the advantage of providing eight wells on each plate where we could add wild-type cells expressing that same GFP-tagged factor, both as a negative control and to monitor possible plate effects.

The library was grown at large-scale and imaged live by three-colour (405 nm, 488 nm, 561nm) automated high-throughput confocal microscopy at high magnification (60x 1.2NA) in 3D *(xy* and 16z planes), following our published protocols^11^. Cells were imaged in blue-fluorescing medium, to be able to visualize them without the need for a cytosolic reporter^11^. In all a total of 1,435,632 images were acquired.

To quantitatively characterize the polarity factors’ localization at growing cell domains, images were computationally analyzed using custom-made image analysis software that: a) segmented cells (blue channel; segmentation was followed by identification of correctly detected cells by Random Forest classifier prediction^11^; b) segmented the RFP-Bgs4 signal (red channel) of each cell to identify growth zones within the cell); c) quantitated within each RFP-Bgs4 growth zone the mean GFP fluorescence intensity of a polarity factor, and then divided it by the mean GFP fluorescence outside the growth zone to obtain its fluorescence enrichment score, for each growth zone in each cell (Figure 1b and c).

**Figure 1.**
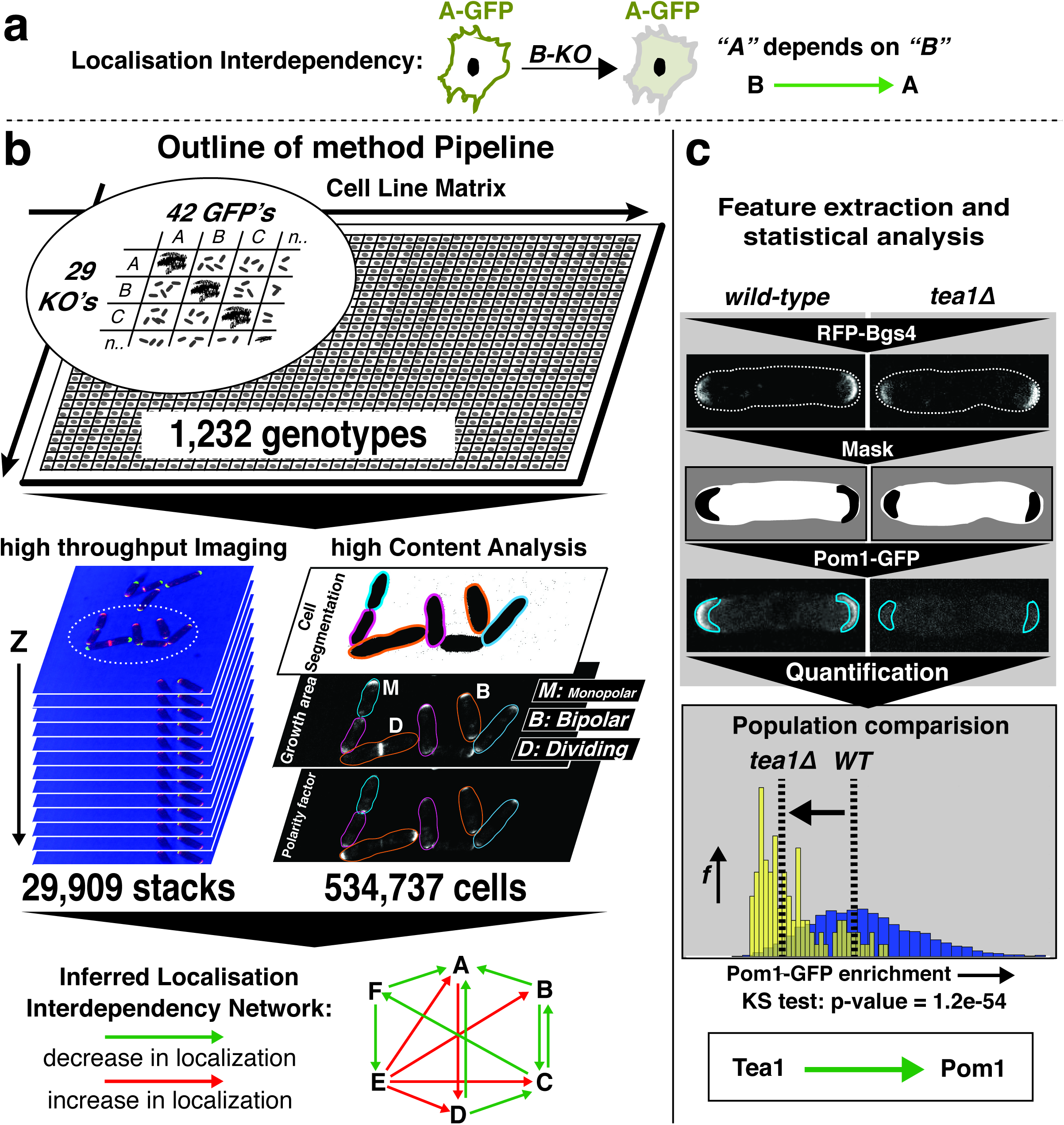
Method for inference of a Localization Interdependency Network (LIN). (a) Definition of a localization dependency. (b) LIN Workflow for the example pathway chosen. A combinatorial library of 1,232 cell lines was generated by crossing 42 cell lines expressing GFP-tagged polarity factors against cell lines knocked out for 29 of those factors. High-throughput, high-resolution imaging of the cell line library followed by computational image analysis was then used to generate high-content image information across separate colour channels. The information was used to assess individual localization dependencies between pairs of factors, and the sum of all significant dependencies allowed construction of the entire interdependency network. (c) Feature extraction and localization dependency detection, exemplified by the change in localisation of Pom1-GFP in a wild-type (WT) versus *tea1* knock-out (*tea1∆*) background^39^. WT and *tea1∆* cell populations co-expressing Pom1-GFP and the RFP-Bgs4 growth reporter were imaged, and areas of cell growth were detected and segmented based on the RFP-Bgs4 signal. This was followed by quantification of the enrichment in Pom1-GFP signal within the growing areas versus the rest of the cell, for all cells. As a result, the distributions of Pom1-GFP enrichment in WT (blue) and *tea1∆* (yellow) cell populations could be compared by KS statistical testing, and a p-value significance attributed to the **Tea1➔ Poml** interaction.

Finally, to establish whether the localization of a given factor **A** depends on another factor **B**, we compared the distribution of *e_GZ_* values of GFP-**A** in **B**-KO versus its distribution in wild-type cells, using non-parametric Kolmogorov-Smirnov (KS) statistics (Figure 1c). If the difference in the distributions was statistically significant as defined by a 1% significance threshold controlling the false discovery rate^18^, A‘s localization was considered to depend on **B**, yielding one oriented edge in the LIN, namely **B➔ A.**

### Quality control

The number of cells used in a pair-wise statistical test is an important consideration, as low numbers can adversely affect statistical sensitivity. While the cell numbers of GFP-tagged polarity factor cell lines in wild-type background were invariably large (n>1200 for all cell lines and n>3500 cells in 80% of cell lines), cell numbers for the individual GFP/KO combination cell lines varied but were adequate (n>100 cells in 80% of cell lines) (Figure 2a and supplementary Figure 2; for a detailed breakdown of cells numbers per cell line, growth stage and of cell length see supplementary table 2). To ensure sufficient coverage as well as assess the reproducibility of our analysis pipeline, we imaged two biologically-independent repeats of each cell line. Computing the correlation of the fluorescence enrichment score *e_GZ_* between the two biological repeats indeed showed a very high reproducibility (Figure 2d), allowing us to conflate the data from both repeats in the subsequent analysis.

**Figure 2.**
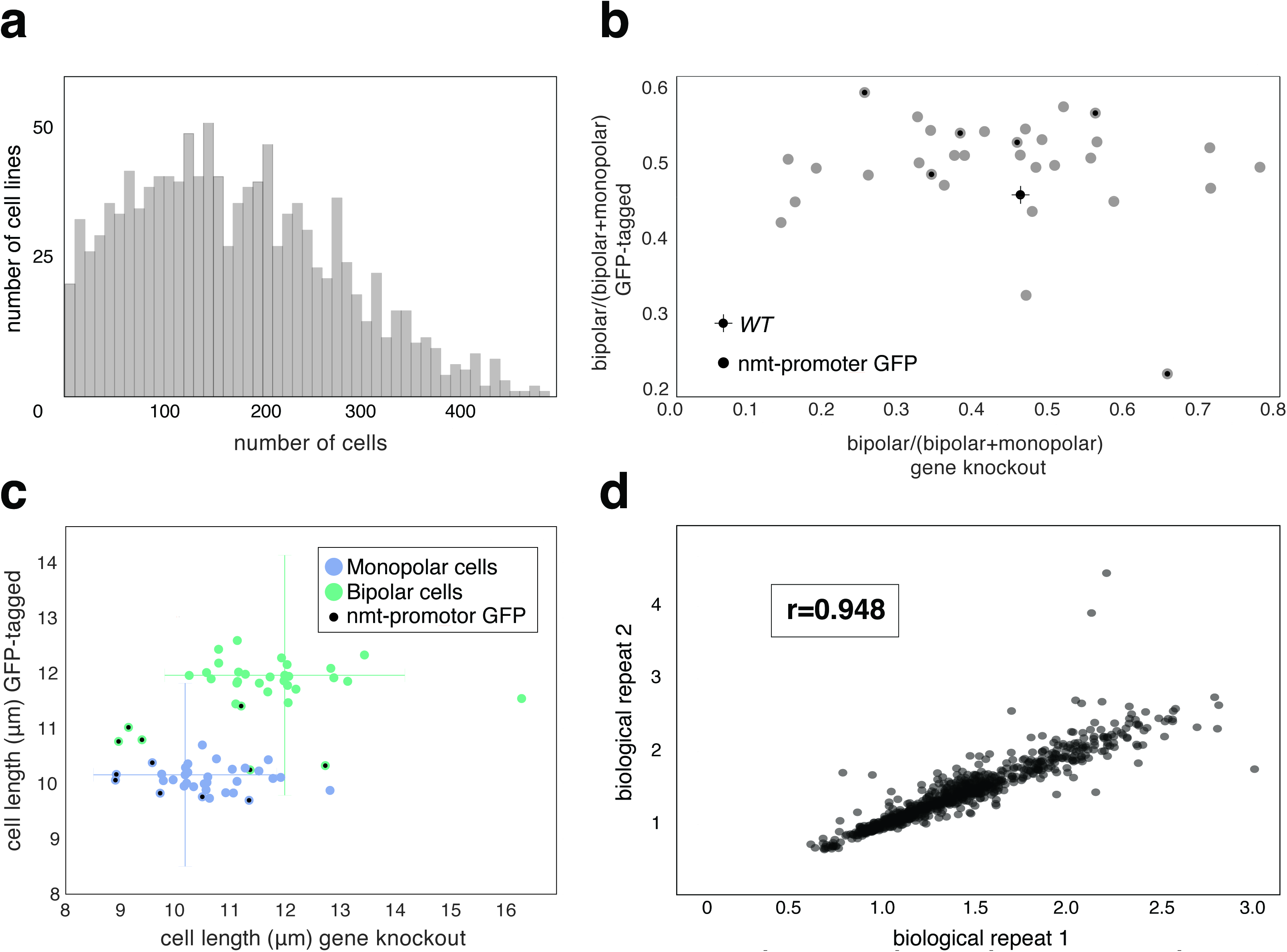
Quality Control. (a) Cell number coverage. Histogram of the total number of cells analyzed across the different GFP/knock-out cell lines. (b) Functionality test of GFP-tags #1. Plot showing the effect on the “bipolar-to-monopolar cells ratio” of knocking-out a polarity factor (x axis) versus GFP-tagging that same factor (y axis). Grey dots: factors expressed under their endogenous promoters. Black dots: factors under nmtl-driven expression. Cross-hair: measurements for the wild-type control (i.e., a cell line without knock-out or tagging; line lengths represent the standard deviations in both axes). (c) Functionality test of GFP-tags #2. Plot showing the effect on average cell length of knocking-out a polarity factor (x axis) versus GFP-tagging that same factor (y axis). Blue/green dots: measurements from monopolar (blue) or bipolar (green) cells. Black dots: factors expressed under nmt1 promoters. Crosshairs: measurements for the wild-type control (i.e., a cell line without knock-out or tagging; line lengths represent the standard deviations in both axes). (d) Reproducibility between biological repeats. Plots of GFP fluorescence enrichment at growing cell domains for all the individual GFP/knock-out combinations, for the two biological repeats of imaging.

Some cell lines however had intrinsically low cell numbers, likely caused by the relatively slower growth and proficiency of their specific gene KOs. This was the case particularly for KOs of the genes *kin1, myo1* and ppbl^19,20^. In those instances further imaging was done to obtain more robust data for statistics.

The biggest potential concern specific to the LIN implementation is the use of GFP-tagging for a large number of proteins, given that any protein-tagging technique has the potential to interfere with gene and/or protein functionality, and induce nonphysiological loss-of-function phenotypes. No previous study has investigated the effect of GFP-tagging at a large scale, prompting us to address it here. Often the functionality of a GFP-construct is individually and qualitatively examined by phenotypic comparison of a cell line expressing it with a non-tagged (wild-type) cell line. Here we chose to systematically and quantitatively compare, for every gene, its KO phenotype versus any phenotype induced by its GFP-tagging, as we reasoned that a likely interference in function by the tag would result in a loss-of-function phenocopy of the KO. We first used as physiological metric the proportion of bipolarly-growing cells relative to all (monopolar and bipolar) growing cells in a population. In the wild-type that proportion is fixed and when disrupted it is indicative of polarity deregulation^21^. As shown in Figure 2b, gene KOs generally disrupted the proportion of bipolar cells (as expected for KOs in polarity regulators^12,13^ while GFP-tagging had a much smaller effect. On average, a gene KO caused a shift in the proportion of bipolar cells that was twice as a large as that caused by GFP tagging (0.125 vs 0.062), indicating the GFP constructs were generally functional. As an additional metric, given the considerable overlap between regulation of polarity and the cell cycle^11,15^ we scored average cell length as a proxy for cell cycle integrity^22^ and found, like before, that gene KOs generally led to much higher disruption of this feature than GFP-tagging, confirming the general functionality of the constructs (Figure 2c). Interestingly, by both metrics the small group of nmt-promoter driven GFP-tagged factors appeared to be less functional than those under endogenous promoter control (Figures 2b and c, black dots), probably because of over-expression effects, vindicating our preference for endogenous promoter-driven expression.

### The LIN and assessment of LIN accuracy

Comparing the distribution of *e_GZ_* values of GFP-**A** in **B**-KO versus its distribution in wild-type cells for all available combinations of GFP-**Y** and **X**-KO yielded the Localization Interdependency Network of 554 edges shown in Figure 3a.

**Figure 3.**
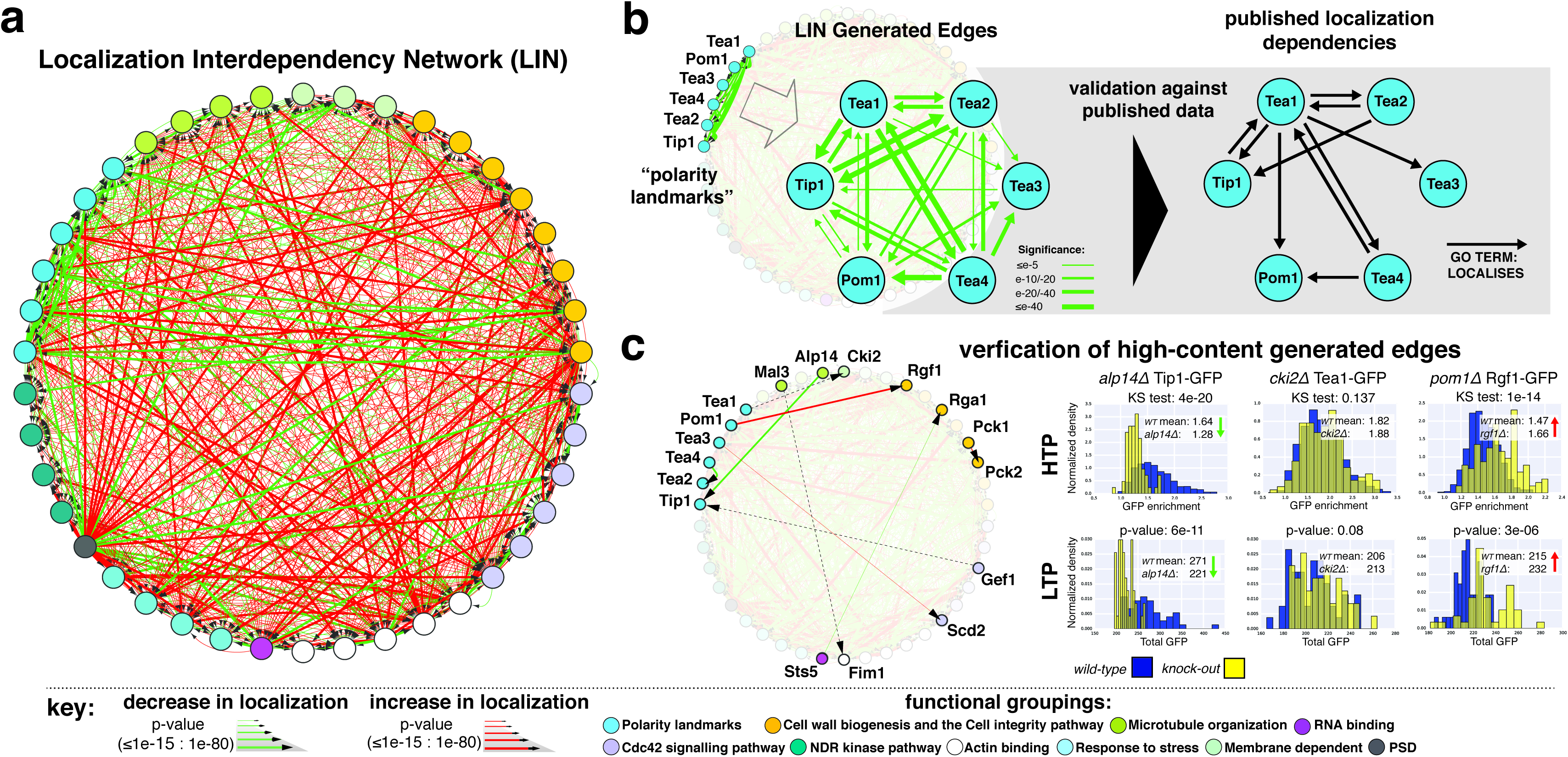
The LIN and assessment of the LIN accuracy. (a) LIN for polarity factors enriched at the fission yeast growing cell domains. Green/red edges signify decrease/increase in fluorescence enrichment at the growing cell domains (edges’ tichkness reflects p-value). (b) Validation against published data. Localization interdependencies amongst the “polarity landmarks” subgroup of polarity factors identified by the LIN, contrasted with those published (annotated under “cellular protein localization” in the Pombase database (http://www.pombase.org/). (c) Verification of high-content generation of edges. Selection of six statistically significant edges within the LIN shown in (a) plus a further three non-significant edges. The high-content generated data (HTP) used to generate the LIN is corroborated well by manually generated (LTP) data for the same three GFP/knock-out combinations (data for the further six GFP/knock-out combinations is included in Supplementary Figure 3).

The 42 polarity factors selected are represented as nodes along the periphery, and have been subdivided into 10 functional groups based on a review of the literature and previously proposed sub-groupings^23^. The edges between the nodes represent 554 statistically significant localization dependencies and are directional from **B➔** signifying ‘KO of **B** affects the localization pattern of GFP-**A**’. The edges’ thicknesses are weighted by the level of statistical significance as defined by the p-value of the KS test, and are colour-labelled red or green to represent a decrease or increase in fluorescence enrichment resulting from a KO, respectively.

From an interpretation point of view, a green edge from B to A represents a cooperative interaction: if absence of B leads to a depletion of A, then B normally enhances the enrichment of A. Conversely, a red edge from B to A represents an antagonistic interaction: B normally suppresses the enrichment of A. Interestingly, out of the 554 significant interactions identified, 334 were found to be antagonistic and 220 cooperative, implying an overall dominance of negative regulatory layers in this pathway.

To probe the accuracy of the inferred LIN, we first checked how many previously published localization dependency interactions for their gene group were present in the LIN. Polarized growth in *S. pombe* has been intensively investigated, and mostly within the functional group of polarity landmarks a number of localization dependencies have been reported in the literature^23^ (the polarity landmarks are microtubule-dependent proteins that mark prospective areas of cell growth^13^. A systematic representation of the published localization dependencies between the 6 polarity landmarks included in our study (annotated under the fission yeast Gene Ontology (GO) search term “localized by”; ^24^ and http://www.pombase.org/) is presented in Figure 3a, alongside a representation of all localization dependencies identified by the LIN. As can be seen in the figure, all 10 known pairwise interactions were correctly identified by the LIN. Additionally, 12 new co-operative interactions were detected, independently confirming the tight functional association between those factors.

Next we verified whether edges detected computationally at high-throughput could be corroborated manually at low-throughput, and as importantly whether absent edges (i.e. edges for which we had enough cells to test them but which were not statistically significant) could also be corroborated. To this end, we selected three randomly-picked novel red edges, three novel green edges and three interactions not deemed statistically significant (Figure 3c). The corresponding cell lines were grown, imaged and analyzed manually using standard low-throughput methods^25^. In agreement with the LIN data, all nine localization dependencies were confirmed by the low-throughput analysis, including their directionality and significance (Figure 3c and Supplementary Figure 3), corroborating the accuracy of the edges detected.

### Biological relevance of the LIN and orthogonal phenotyping

For the LIN method to be of biological relevance, the collection of statistically significant localization dependencies it identifies should provide new biological insights that can be further tested and validated, including by orthogonal (non-image based) phenotyping.

To probe the discovery power of the LIN method in our specific example pathway, we sought to validate a prominent and unexpected pattern of localization dependencies we observed between two polarity factor subgroups: the polarity landmarks (shown in Figure 3b) and the components of the cell integrity pathway (CIP; ^26^), a Rho-GTPase/protein kinase C dependent pathway that co-ordinates different cellular functions including cell wall growth and that in response to external stresses activates the MAP-kinase Pmk1 through a conserved MAP-kinase cascade ^26^. Specifically, we found a consistent reduction in fluorescence enrichment of the polarity landmarks Tea1, Tea4 and Pom1 in KOs of the CIP members Rgf1, Pck1, Pck2 and Pmk1, suggesting that the CIP members generally cooperate in localizing the polarity landmarks at the cell cortex (Figure 4a). Conversely and interestingly, we found a consistent increase in localization of those CIP members in KOs of the polarity landmarks, suggesting that the landmarks generally antagonize the localization of the CIP components (Figure 4a). No evidence of bi-directional, pathway-wide interaction between the CIP and the polarity landmarks exists in the literature, making their link inferred by quantitative imaging a good candidate for orthogonal validation.

**Figure 4.**
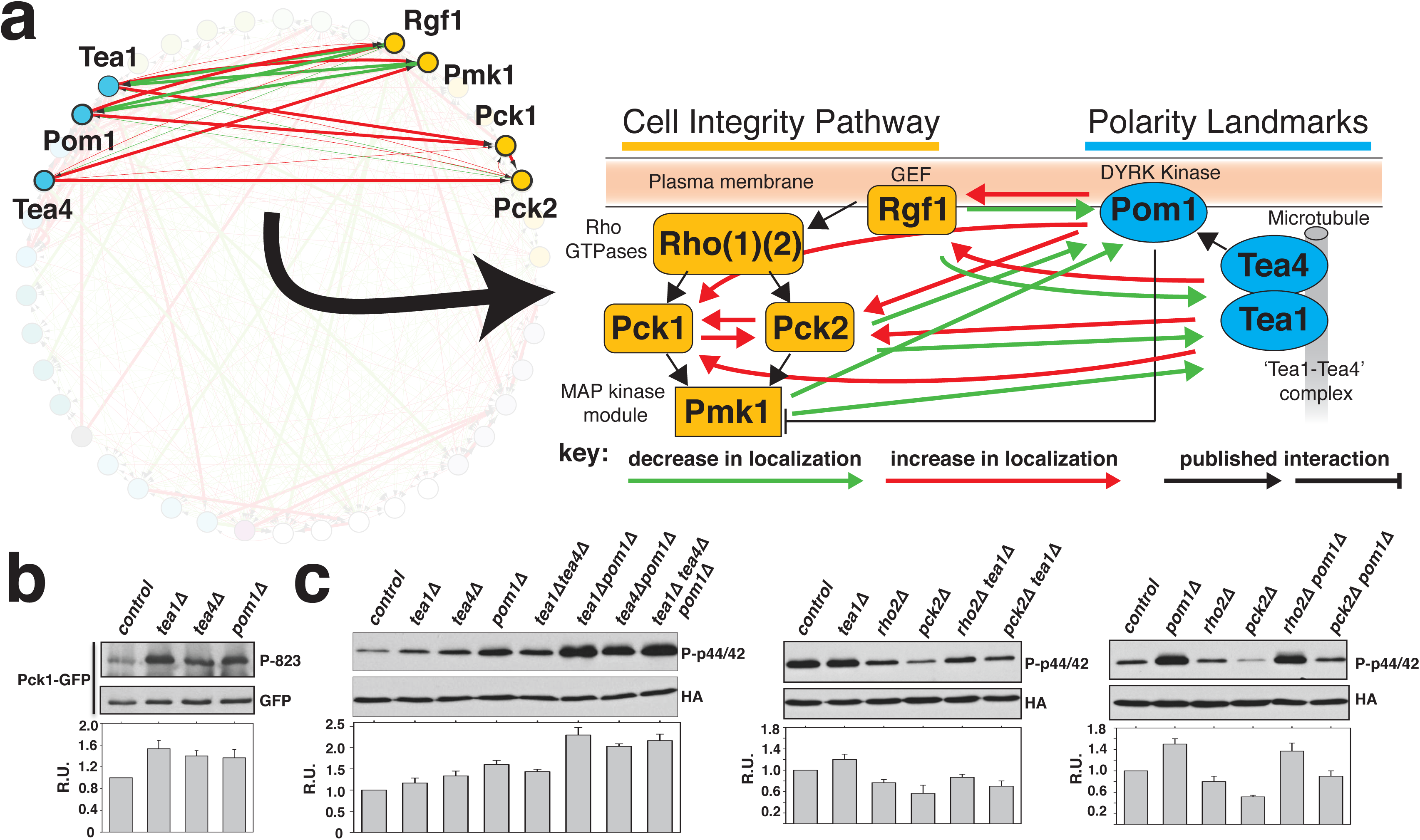
LIN method validation by orthogonal phenotyping. (a) Systematic localization interdependencies observed between the polarity landmark factors Tea1, Tea4 and Pom1 and the cell integrity pathway (CIP) components Pck1, Pck2, Pmk1 and Rgf1, represented on the LIN (left) and overlaid to published, known interactions (right). (b) The level of Pck1 activation loop phosphorylation at T823 (and hence activity) in polarity landmark knock-outs. The interaction inferred from this method matches the LIN inferred interaction. (c) Left panel: Pmk1 phosphorylation (and hence activity) level changes in single, double and triple polarity landmark knockouts. This is dependent on both *pck2* and *pck1* (supplementary Figure 4c) in agreement with the LIN predictions. (NOTE: “RU” is relative units standardized to the control. All phosphorylation experiments were repeated three times with similar results).

Given that the CIP members Rgf1 (Rho-GEF) and two PKC-orthologues Pck1 and Pck2 are involved in cell wall synthesis and remodelling at the cell ends, we reasoned their relative enrichment there could reflect an increased activation of the pathway by phosphorylation, which we should be able to detect biochemically. In agreement with this, we observed that specific activation loop phosphorylation of Pck1 at T823 was increased in the absence of the polarity landmarks (Figure 4b and supplementary Figure 4a). To probe this further and given that the CIP activity is reflected in the phosphorylation status of Pmk1^27^, we also determined its phosphorylation level in control versus single, double, and triple mutants lacking different combinations of the polarity landmarks during growth and in response to stress (Figure 4c and supplementary Figure 4b,). Strikingly, we found that knockout of the polarity landmarks led to increased basal activity of the CIP and activation upon salt stress dependent on both Pck2 and Pck1 (supplementary Figure 4c, d), consistent with the predictions of the LIN (Figure 4a). Interestingly, the increase in Pmk1 phosphylation may occur through separate mechanisms, since Pmk1 phosphorylation in *tea1/tea4* mutants was additive to that of *pom1* KO (Figure 4c), and was mostly Pck2-dependent in the case of the *tea1/4* KO, but only partially so in the *pom1* KO (supplementary Figure 4c)^28^. Pck1 might be responsible for this alternative regulation, since its knockout lowered Pmk1 phosphorylation in the polarity landmarks knock-outs, and this effect was more pronounced in the *pom1* KO (supplementary Figure 4d).

Loss of polarity factors might somehow elicit a distortion at the plasma membrane and/or cell wall to elicit a downstream activation of the CIP. In the distantly related budding yeast *Saccharomyces cerevisiae* the CIP has been found to regulate membrane lipid homeostasis^29^. Therefore, decreased polarity factor retention in CIP knockouts might also reflect a reduction in the integrity of the plasma membrane. These biological findings will need to be further investigated in the future.

## DISCUSSION

In short, the LIN method provides a powerful tool to generate statistically weighted and cross-comparable relational information that can be used to accurately predict pathway information. The strengths of the LIN method for biological pathway discovery come from the fact that functional interactions are inferred from high-content, spatio-temporally resolved information provided by the single-cell imaging, and that interactions’ directionalities are directly detected, rather than mathematically inferred for example by Bayesian network analysis^9,11,30,31^. As a consequence of this, contrary for example to the undirected relationships obtained by mapping protein-protein or synthetic genetic interactions^32^, the LIN method allows inference of networks of systematically assessed, statistically-weighted and intrinsically directed interactions. In addition, the spatio-temporal information assayed within individual cells likely provides stronger sensitivity than other population-based methods as it can detect interactions that are relatively subtle, compared for example to the gross changes in cellular fitness scored in Synthetic Gene Arrays ^4,33^

In the example case chosen, we found the cell polarity LIN to be extensive and accurate, and that it both recapitulates available knowledge and provides novel and subtle, systems-level biological insights. However, the example shown also underlines some of the general challenges of building LINs.

First, the absence of edges needs to be interpreted with caution, given that some edges might be inaccessible simply due to experimental constraints, like low cell numbers obtained for specific GFP/KO combinations (which compromises statistical sensitivity) or the fact that some combinations might be non-viable (for example, due to some of the genes disrupted being essential or to specific GFP/KO combinations being synthetically deleterious). Whilst low cell numbers can be improved by additional imaging, non-viable GFP/KO combinations represent a structural problem that cannot be overcome and intrinsically sets an upper limit on the completeness of inferred LINs.

Secondly, by construction the physiological accuracy of LIN is vulnerable to loss of protein functionality due to gene tagging. Specifically in the example shown, the phenotypic defects found in GFP-tagged cell lines were in general weak with the exception of cell lines using non-endogenous promoters, which likely reflect nonphysiological levels of transcription. This highlights the need with the LIN approach to assess with caution the functionality of each individually tagged cell line/construct to be analyzed, and the selective-removal of conditions suggestive of loss of function. Thirdly, localization dependencies may reflect indirect downstream or cross-system interactions between factors, rather than direct protein-protein mediated interactions, potentially obscuring interpretation of the information contained in the LINs. A challenge for future work will be to disentangle the causality of direct versus indirect interactions inferred by this approach, either experimentally using biochemical techniques or mathematically using techniques such as signed transitive reduction^34^ or other network simplification approaches^35^.

As described and out of simplicity, this study focused on only one image-based feature and phenotype to build one LIN, using data from yeast. However, the method can be in principle implemented using any species and cellular system of choice, and can be extended to encompass multiple quantitative features and other phenotypes, including time-resolved phenotypes^36,37^, allowing to generally harness the full breadth of information available in image-based screens. Based on that information cells within a population could be sub-pooled according to specific characteristics, such as morphology, cell cycle stage or other, allowing deconvolution of the LIN into multiple sub-networks reflective of specific physiological states. Cross-comparison of those sub-networks could indicate functional roles for defined pathway features in mediating the transition from one physiological state to another. Likewise the approach could be extended to examine the LIN and selected physiological states in the presence of insults or drugs^38^, allowing insights into the contextual re-wiring of the regulatory networks in these environments.

## METHODS

Methods and any associated references are available in the online version of the paper.

Note: Supplementary information is also available in the online version of the paper.

## ACKNOWLEDGMENTS

We thank J. Bahler, H. Moriya, K. Nakano, M. Toya, J. Pines, M. Godinho Ferreira, T. Surrey, J. Howard, A. Ciliberto, C. Pal, A. Sveiczer, C. Bakal and the Carazo-Salas, Csikász-Nagy and Sato groups for help and comments, and J. Bahler and C. Bakal for critical reading of the manuscript. This work was supported by a Human Frontier Science Program (HFSP) Young Investigator Grant (R.E.C.S., A.C.N., M.S.; HFSP RGY0066/2009-C), an European Research Council (ERC) Starting Researcher Investigator Grant (R.E.C.S.; SYSGRO), a Biological Sciences Research Council (BBSRC) Responsive Mode grant (R.E.C.S.; BB/K006320/1), two Isaac Newton Trust research grants (R.E.C.S.; 10.44(n) and 14:39(h)), a CRUK-funded Cambridge Cancer Centre (CCC) Pump Priming Award (R.E.C.S.), startup funds from the University of Bristol (R.E.C.S.), the Italian Research Fund FIRB (A.C.-N; RBPR0523C3), a Grant-in-Aid for Young Scientists (A) from the Japan Society for the Promotion of Science (JSPS) (M.S.) and grants from the Ministerio de Economía y Competitividad (J.C.; MINECO BFU2011-22517 and BFU2014-52828-P).

## AUTHOR CONTRIBUTIONS

R.E.C.-S., A.C.N. and M.S. conceived/led the project and designed the general experimental and computational strategy. M.S., M.Y. and K.A. generated the GFP-tagged cell lines. J.D. generated the combinatorial GFP/KO library, and carried out all microscopy phenotyping experiments, with help from M. Geymonat and J.F.A.. A.C. developed the image processing and large-scale analysis pipeline. F.V. carried out thorough statistical quality control with help from M. Giordan. M.M. and J.C. carried out biochemical, orthogonal phenotyping experiments. J.D., A.C. and R.E.C.-S. wrote the text with help from F.V. A.C.N., and other co-authors.

## Supplementary Figure Legends

**Figure S1.**
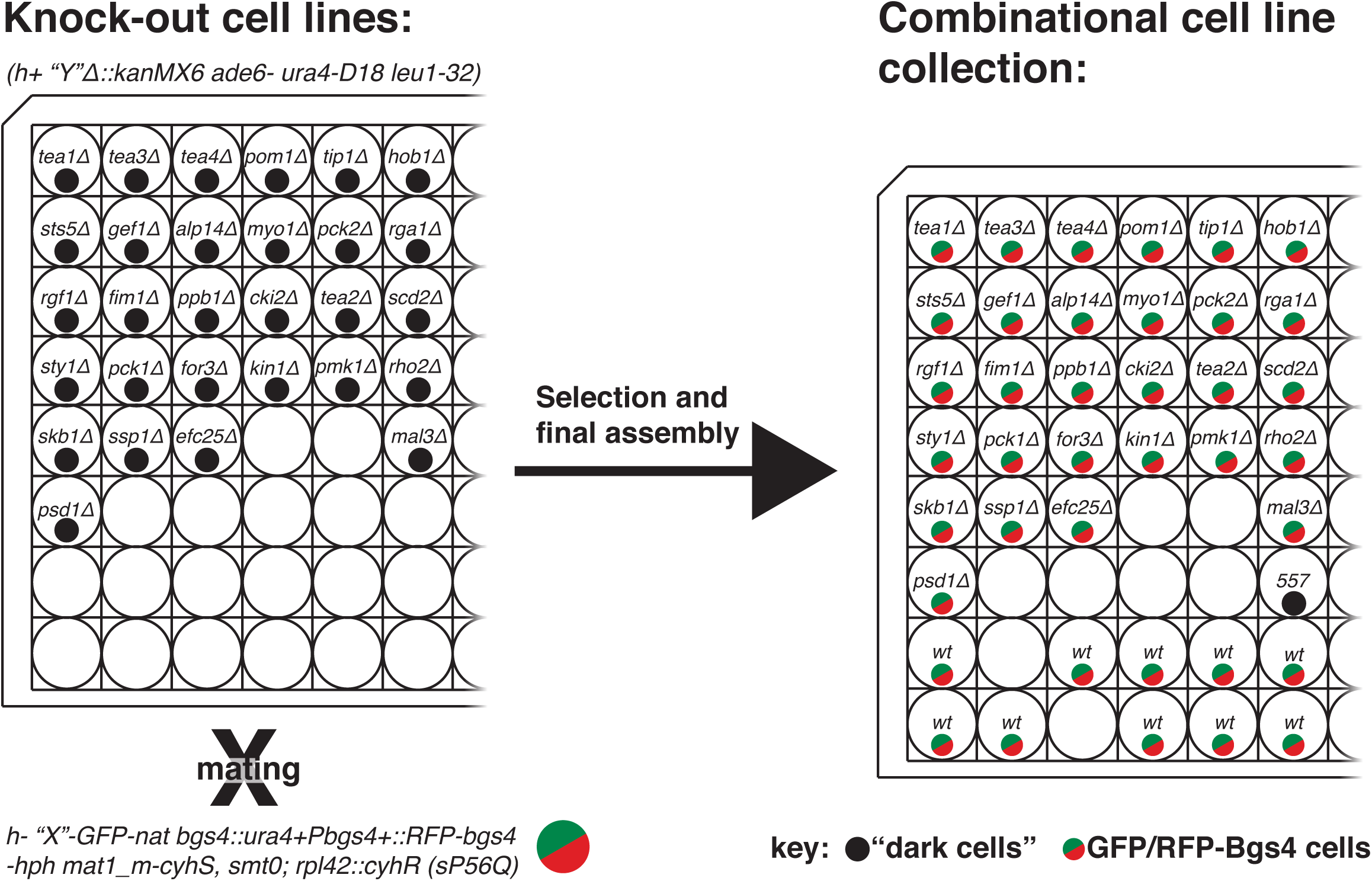
Construction and physical layout of the cell line library. The cell line library was principally constructed by mating the collection of knock-out cell lines with the individually GFP-tagged cell lines, followed by drug selection of the appropriate progeny. The final library was completed by the addition of the same GFP-tagged cell line in a WT background and a “dark” cell line.

**Figure S2.**
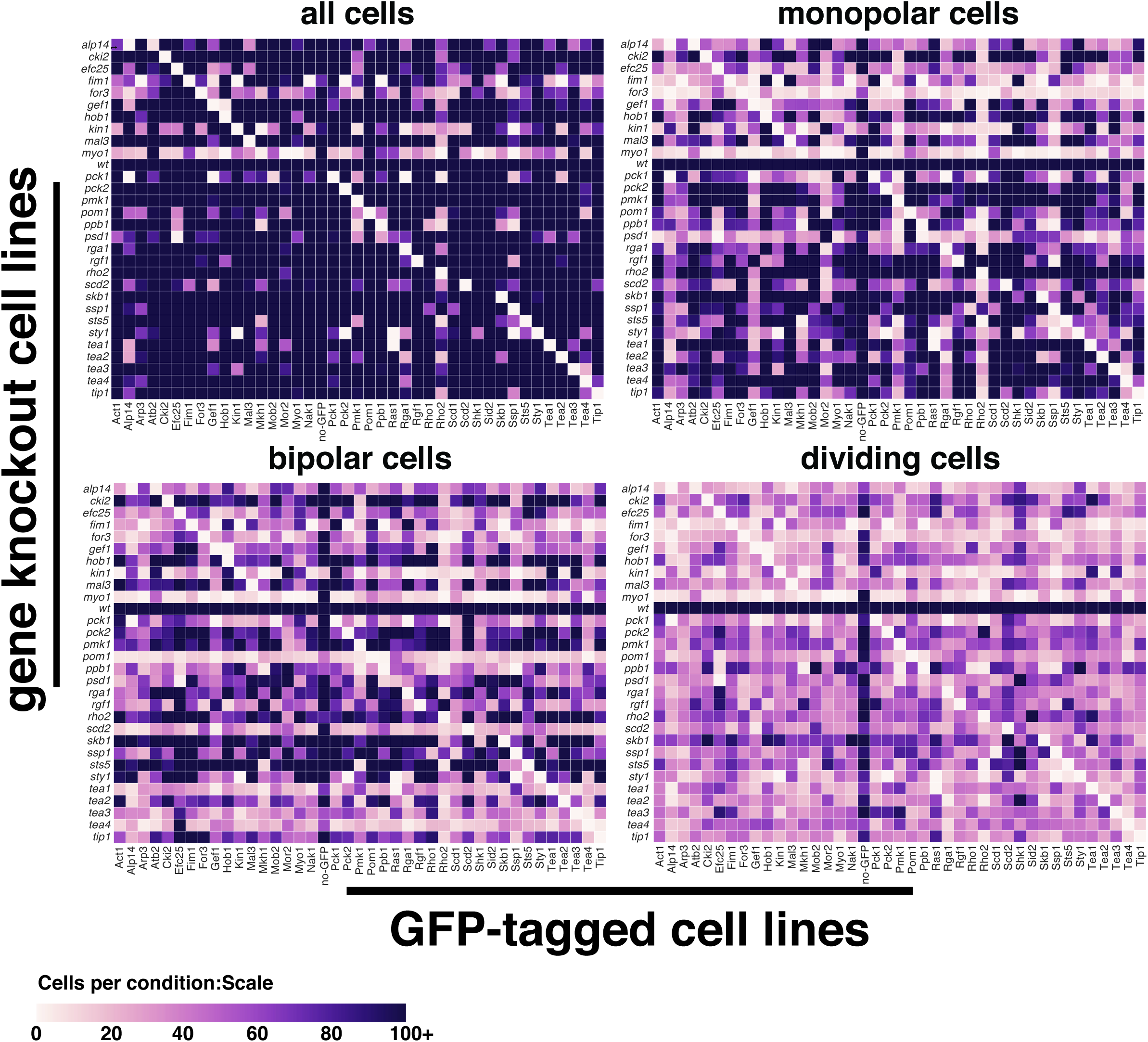
Cell numbers for the individual GFP/KO combination and WT cell lines. Matrices are divided into cells growing monopolar, bipolar, dividing and all cells.

**Figure S3.**
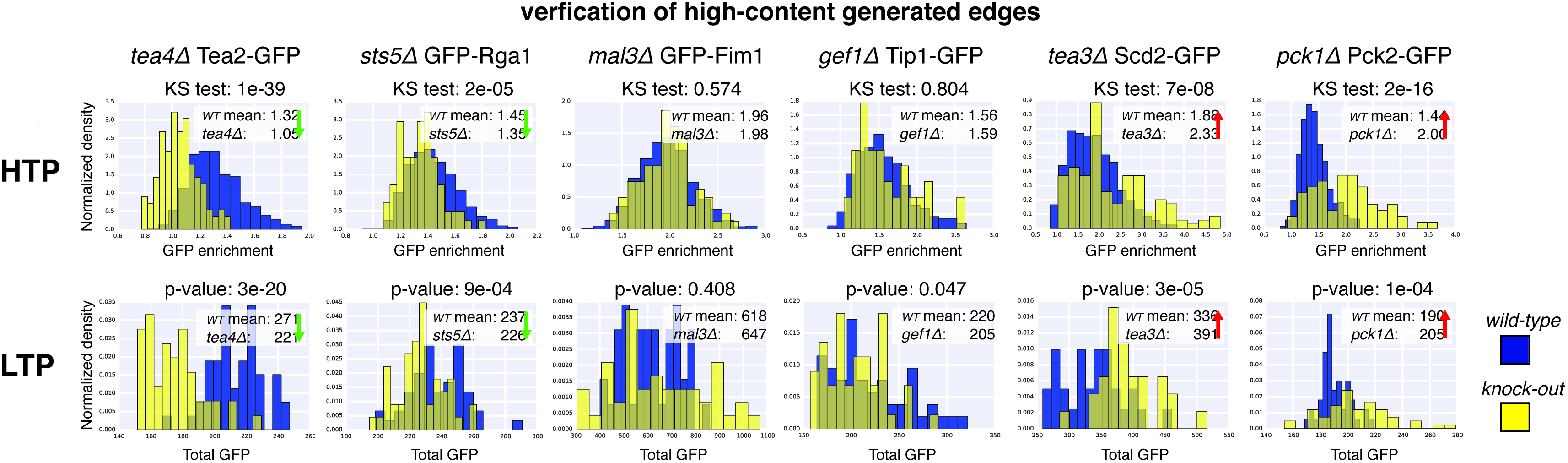
Verification of high-content generation of edges. The high-content generated data (HTP) used to generate the LIN is corroborated well-by-well, by manually generated (LTP) data for a further six GFP/knock-out combinations listed in Figure 3.

**Figure S4.**
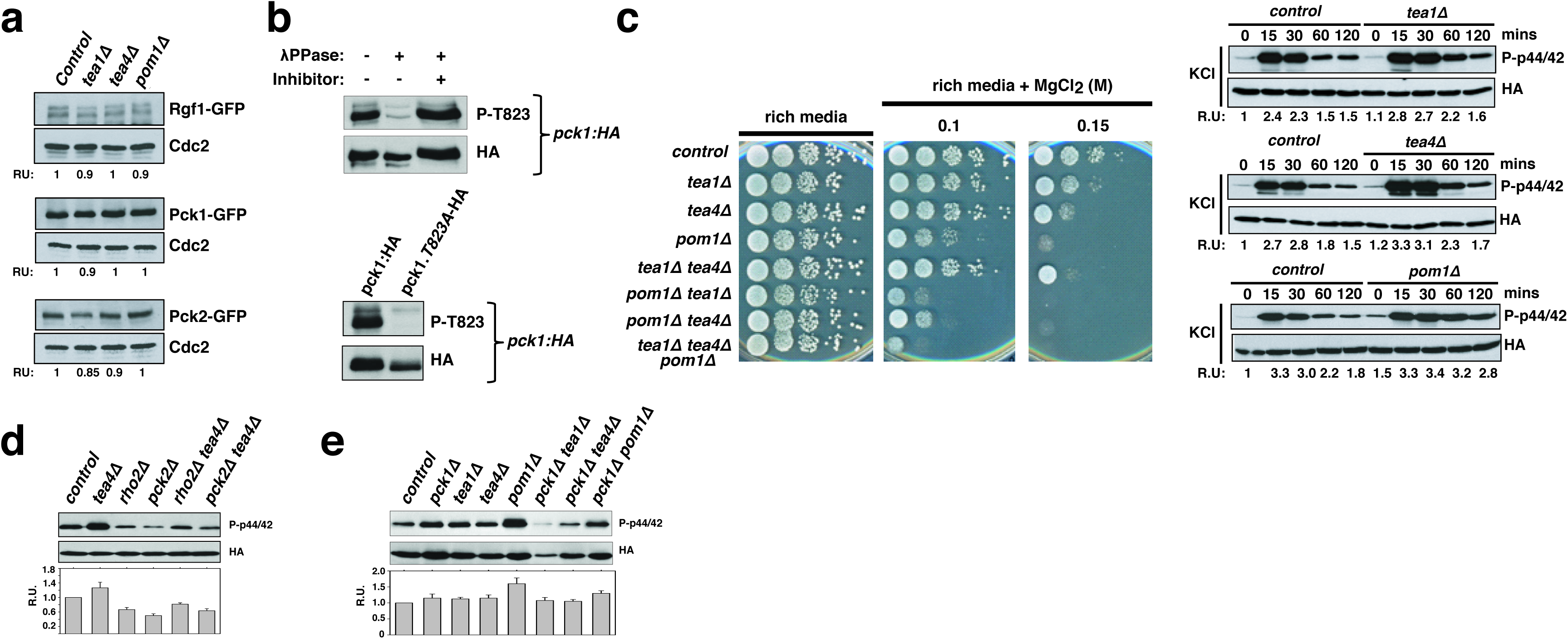
LIN method validation by orthogonal phenotyping. (a) Demonstration of the specificity of the anti-phospho T823 antibody. (b) Serial-dilutions of single, double and triple polarity landmark knock-outs exhibit hyper-sensitivity to MgCl_2_ a phenotype associated with activation of the CIP. Pmk1 phosphorylation level changes in response to potassium chloride (KCl) are altered in the polarity landmark knockouts. (c) Pmk1 phosphorylation in the *tea1* and *tea4* knock-outs is mostly dependent on Pck2, but in the *pom1* knock-out is only partially dependent on Pck2. (d) Increased Pmk1 phosphorylation in the *tea1, tea4* and *pom1* knock-outs is lowered by further knockout of Pck1.

## ONLINE METHODS

### Cell lines and combinatorial library construction

All drug selection was performed on solid yeast extract with supplements (YES)(Moreno *et al.* 1991) with concentrations of G418 at 200 mg/L, nourseothricin (Nat) at 100mg/L, hygromycin (Hph) at 100mg/L and cycloheximide (CycH) at 100mg/L. Mating was performed on solid EMM lacking nitrogen (EMM-N) (Moreno *et al.* 1991). All incubation steps were performed at 30^0^C.

The GFP-tagged cell lines and knockout cell lines used to construct the cell line combinatorial library are listed in supplementary table 1. Most (37 out of 42 total) of the GFP constructs were created for this project (see the “Molecular Genetics” section) the other cell lines were sourced from other labs. All GFP cell lines constructed specifically for this project were qualitatively assessed for signal and functionality. As a satisfactory GFP tag of Psd1 could not be constructed it was finally decided not include this in the final library of cell lines. The knockout cell lines were mainly sourced from the Bioneer haploid deletion library version 2 and 3 (Bioneer, S. Korea), others were constructed in particular for this project (see “Molecular Genetics section”) and also sourced from other labs. All the Bioneer derived knockout cell lines were verified by PCR testing.

To remove diploids resulting from the mating process and ensure the progeny are exclusively of a single mating-type (h^+^) we utilized the PEM-2 system (Roguev *et al.* 2007). The GFP-tagged constructs were converted into a *mat1_m-cyhS, smt0; rpl42::cyhR (sP56Q) RFP-Bgs4* background (Table 1) through a novel mating and selection procedure. Firstly the GFP-tagged cell lines were mated with an *RFP-Bgs4-hph* cell line (cell lines M54 and M57 a kind gift from Marco Geymonat) and appropriate h^+^ progeny selected. These were then mated with the PEM-2 P392 master cell line and the progeny were selected on YES plates containing Nat and Hph (vegetative cells were removed from the mating mixture by incubation in 40% ethanol for 20 minutes). To establish if individual colonies had the background *mat1_m-cyhS, smt0; rpl42::cyhR (sP56Q),* firstly CycH sensitive colonies were identified and then mated with a *h+ Tea1∆::KanMX6* cell line and tested for the ability to generate viable colonies on media containing CycH, and G418 (CycH sensitive parent cell lines that can generate CycH resistant progeny, have the background *mat1_m-cyhS, smt0; rpl42::cyhR (sP56Q)).*

The process of construction and physical layout of the cell line combinatorial library is summarized in Figure S1. A high-through-put mating strategy for construction of the combinatorial library made use of a manual 96-well pipettor (Rainin Liquidator 96 Manual 96-well Pipettor, Mettler-Toledo, Switzerland). A small volume of liquid culture (~2μl) containing a mix of the corresponding cell line on the knockout array and the PEM-2 “X”-GFP for that half-plate was spotted onto an omni-tray (Thermo Scientific, UK) containing EMM-N. Following an eight day incubation the mating mixture at each position was pinned onto plates containing YES+CycH incubated for two days and then pinned onto selective YES+CycH,G418,Nat,Hph plates. The final plates were created by addition of the corresponding WT “X-GFP” cell line. Cells were frozen for long-term storage at -80°C in 50% yellow freezing mix (YFM, 50% glycerol).

Relatively few cells were segmented for cell lines with the background *Kin1∆, Myo1∆* and *Ppb1∆.* Consequently further plates were constructed and imaged with these knockouts crossed to a GFP array containing the 32 endogenously expressed GFP cell lines.

### Molecular Genetics

The constructs created specifically for this project were made using the following methods:

Twenty-one GFP C-terminally tagged cell lines were created using standard methods^40-42^. Namely, DNA fragments containing genes encoding GFP or 3GFP (three tandem copies of GFP) and clonNAT resistance marker (nat) were prepared by PCR with the template plasmid pMS-GFP-nat. The fragments were then introduced into WT cells (JX92, *h ^−^his7 leu1-32 ura4-D18 ade6-M216*) to be integrated into the C-terminal end of a target gene. Stable transformants were selected in YE5S containing cloNAT (100 μg/ml). The C-terminal end of each protein and the N-terminal head of GFP (or 3GFP) were connected with a linker (amino acid sequence: GAGAGAGAGAFSVPITT).

Six cell lines expressing N-terminally tagged GFP-fusion proteins expressed form their endogenous promoters were created. The GFP-Atb2 cell line has been previously described^43^, the remaining six cell lines were made as follows: first a intermediate *ura4*^+^-Pnmt1-GFP-‘X’ cell line was made, through the standard PCR-based method using the plasmid pNGU (pFA6a-*ura4*^+^-Pnmt1-GFP-linker) as a template. The *ura4^+^-* Pnmt1 region was then excised out by the DNA fragment connecting the upstream (promoter) region of *the target gene* and GFP sequence, and cells in which the excision occurred correctly were selected on YE5S plates containing 5-fluoroorotic acid (1 mg/ml). As a result, the promoter region of *the target gene* was directly connected to the GFP coding sequence to express the GFP-‘X ‘fusion protein under the native promoter. Finally, the natMX selection marker was inserted at the upstream region of the target genes promoter.

N-terminal GFP-fusion genes of another nine cell lines were expressed by the *nmt1* promoter (Pnmt1) or its weaker variant (Pnmt41). These were constructed using plasmids pFA6a-kanMX6-p3nmt1-GFP or pFA6a-kanMX6-P41nmt1-GFP as templates for PCR, respectively. After constructing integrants conferring resistance to G418, the kanMX marker was replaced with the natMX marker through the marker switch method^41^.

For actin visualization, Lifeact^44^ was fused to the N-terminus of GFP (LA-GFP) and expressed under the *dis2^+^* promoter. The DNA fragment containing or Pdis2-LA-GFP-nat was inserted into the co2 region (at the base number 2939887-2939890) of Chromosome 1 of a WT cell line.

### High throughput preparation and plating of the cell line combinatorial library for imaging

The high throughput growth and plating of the cell line combinatorial library was based on the protocol developed and extensively recorded by Graml et al. 2014. Briefly the cell lines were grown in 96-well plates with liquid YES broth (or for those cell lines with nmt-promotor driven GFP constructs; in EMM liquid) for over 48hours at room-temperature. The large-scale synchronization of growth state between the multiple cell lines was achieved through a system of serial dilutions. For imaging, cells were adhered to glass-bottomed microwell plates (Swissci, Switzerland) precoated with lectin (EY Laboratories, USA), and lastly suspended with 100 μ1 of liquid YES or EMM medium plus 0.1 mg/ml Dextran Cascade Blue 10.000 MW (Invitrogen, USA).

### High throughput imaging of the cell line combinatorial library

Imaging was performed on the spinning-disc confocal OperaLX high-throughput microscope system (PerkinElmer, USA) with a 1.2 NA 60x water immersion objective. This microscope system is fully automated and combines robust auto-focusing with high-powered immersion optics. Settings used for imaging were: z-stack (16 planes, 0.4 μm separation), using exposure channels blue (405nm, 40 ms, full power (~1000 μW)), green (488 nm, 40 ms or 160 ms, full power (~5000 μW)) and red (561nm, 40 ms, full power (~2,400 μW). Imaging of each 96-well plate comprised imaging a field in each well then five additional runs in a new field (the field positions were spread throughout the well).

### Image Analysis

Single cell segmentation was performed as described in Graml and al. 2014^11^, using the blue channel where the cells appear as dark on a bright background thanks to the addition of Dextran Blue in the medium. Within each cell, the red channel was segmented by thresholding to provide a mask of the growth area of the cell.. For Fig. 2B, growth stage (monopolar or bipolar) was automatically assigned by looking at the number and location of growth areas in each cell.

Analysis of the green channel proved more challenging as the tagged protein had various localisation within the cell in wild-type background and the KO could lead to fairly diverse phenotypes. A more thorough look into those phenotypes will be described in a forthcoming study. Here, to present LINs and demonstrate their power as a tool to reconstruct pathways at a system level with a spatio-temporal resolution, we focus on one feature, the fold enrichment in fluorescence of the GFP tagged protein in the growth area with respect of the rest of the cell. Most proteins of interest are known to be enriched in the growth areas one way or another and this is the single most informative feature we could compute. As with any quantitative fluorescence measurement, we had to be very careful in the preprocessing of the image. The camera background level was subtracted from the image, and the uneven illumination of the microscope was corrected, with the illumination pattern being estimated on a plate by plate basis using the medium autofluorescence in an empty well.

### Quality Control

As a quality control we compared the effect of GFP tagging with the effect of gene knockout on two distinct phenotypes: the mean cell length, and the bipolar-to-monopolar cell ratio. Specifically, we computed the average cell length (or, the fraction of bipolar to monopolar cells) for every wild-type cell line expressing a given GFP-tagged polarity factor, and compared it to the cell length of the corresponding ‘dark’ (non-GFP tagged) cell line knocked out for that polarity factor. For example, the Tea1-GFP cell line was compared with the untagged *tea1∆* cell line. The reference cell line shown in the middle of the plot (with a cross) corresponds to wild-type fission yeast, without any gene GFP-tagged or knocked out. As we can see from the plot, knockouts tend to cause a much bigger change in the phenotype than GFP-tagging. Computing the correlation of the fluorescence enrichment score *e_GZ_* between the two biological repeats was created by simply measuring the mean of our feature of interest separately for cells from each of the two repeats done for each cell line and computing the correlation.

### Network Inference

Using the imaging data, we constructed a signed, directed and weighted network among all cell end-localized proteins in our screen.

We used the nonparametric Kolmogorov-Smirnov Test to compare the GFP cell end enrichment in the background of a given knockout with the cell end enrichment in the corresponding wild-type cell line. To calculate the KS test, we used the implementation present in SciPy, an open source highly mature scientific software library (www.scipy.org).

If the test was significant, an edge between the knockout and the GFP tagged protein was drawn. For example, to test whether Pom1 localization is controlled by Tea1, we compared the Pom1 cell end enrichment in *pom1-GFP-WT* vs *pom1-GFP-tea1∆* cell lines.

To bound the number of false positives due to the multiple testing we choose to control the false discovery rate with the method of Benjamini-Hochberg (1995)^18^. A test was considered significant if its adjusted p-value was below 0.01. P-value correction was carried out using statsmodels (http://statsmodels.sourceforge.net/) an open-source statistical toolbox for Python.

To estimate the magnitude of the effect (the weight of the edge in the network), different choices are possible. A negative log-pvalue is a natural candidate but with a huge sample size can be really large also if the magnitude of the effect is small. We used the value of the KS-statistic, which corresponds to the maximum difference between the two empirical distribution functions. This statistic can be influenced both by location and shape differences of the two distributions and does not have the previous drawback.

The sign of the interaction was determined by the direction of the change of the cell end enrichment. If a protein’s cell end enrichment was decreased upon knockout of another, we called the interaction positive. If the protein’s cell end enrichment increased, the interaction was negative.

### Low-throughput verification of the high-throughput data

Selected cell lines for low throughput verification were grown in liquid YES to exponential growth phase and plated on lectin pre-coated glass-bottomed dishes Matek, USA) and finally suspended in YES. Imaging was performed with a DeltaVision (Applied Precision), comprising an Olympus 1X71 widefield microscope, an Olympus UPlanSapo x100 oil immersion lens (NA1.4) and a Photometrics CoolSNAP HQ2 camera. Stacks of 16 planes 0.4μm apart with GFP, RFP and transmission channels were taken (with FITC, TRITC and POL filter sets respectively). Stacks were captured, processed and deconvolved using SoftWoRx (Applied Precision).

Low throughput image analysis was performed manually in Fiji with the deconvolved stacks. Selected cell-ends with the appropriate growth phase (as determined from the RFP-Bgs4 signal) were identified and uniformly re-orientated and positioned at their cell-ends. A single constant region of interest (ROI) following the approximate contours of the cell-end was positioned accurately at the cell-end apex using the RFP-Bgs4 signal for guidance. The average pixel intensity within the ROI for the GFP channel was measured and recorded.

### Biochemical Techniques

*Detection of total and activated Pmk1-* Cells from log-phase cultures (OD_600_= 0.4) of either control or mutants expressing a genomic Pmk1-HA6his tagged fusion and growing in YES medium were collected during unperturbed growth or after treatment with 0.6M KCl (Sigma Chemical). In Pck1 overexpression experiments plasmid pREP41X-*pck1*^+^ was used^45^, and transformants were grown in EMM2 minimal medium for 24 h with or without 5 g/ml thiamine (Sigma Chemical). Preparation of cell extracts and purification of HA-tagged Pmk1 with Ni^2+^-NTA-agarose beads (Qiagen) was performed as previously described^27^. The purified proteins were resolved in 10% SDS-PAGE gels and transferred to Hybond-ECL membranes (GE-Healthcare). Dually phosphorylated and total Pmk1 were detected with rabbit polyclonal anti-phospho-p44/42 (Cell Signaling) and mouse monoclonal anti-HA antibody (12CA5, Roche Molecular Biochemicals), respectively. The immunoreactive bands were revealed with an anti-mouse-HRP-conjugated secondary antibody (Sigma) and the ECL system (Amersham-Pharmacia). Densitometric quantification of Western blot signals was performed using Image J^46^. Experiments were repeated at least three times with similar results. Relative Units (RU) for Pmk1 activation are estimated in each experiment by determining the signal ratio (as a measurement of band intensity) of the anti-phospho-P44/42 blot (activated Pmk1) with respect to the anti-HA blot (total Pmk1) at each time point.

*Detection of total Rgf1, Pck1, and Pck2-* Cell extracts were prepared under native conditions using Buffer IP (50 mM Tris-HCl (pH 7.5), 5 mM EDTA, 150 mM NaCl, 1 mM b-mercaptoethanol, 10% glycerol, 0.1 mM sodium orthovanadate, 1% Triton X-100, protease inhibitors at standard concentration). Equal amounts of total protein were resolved in 6% SDS-PAGE gels and transferred to Hybond-ECL membranes. Total Rgf1-GFP, Pck1-GFP, and Pck2-GFP were detected with polyclonal rabbit anti-GFP (Cell Signalling) antibody. Anti-PSTAIR (anti-Cdc2, Sigma Chemical) was used for loading control.

*Detection of phosphorylated Pck1-* To detect *in vivo* phosphorylation of Pck1 at T823, an anti-phospho-polyclonal antibody was produced by immunization of rabbits with a synthetic phospho-peptide corresponding to residues surrounding Thr823 of Pck1 (GenScript).

*Reproducibility of results* Densitometric quantification of Western blot signals was performed using Image J ^46^. Experiments were repeated at least three times with similar results. Mean relative units ± SD and/or representative results are shown.

### Plate assay of stress sensitivity for growth

Appropriate decimal dilutions of *S. pombe* control and mutant cell lines were spotted per duplicate on YES solid medium or in the same medium supplemented with different concentrations of MgCl_2_ (Sigma Chemical). Plates were incubated at 28°C for 3 days and then photographed.

---

**Supplementary table 1.**
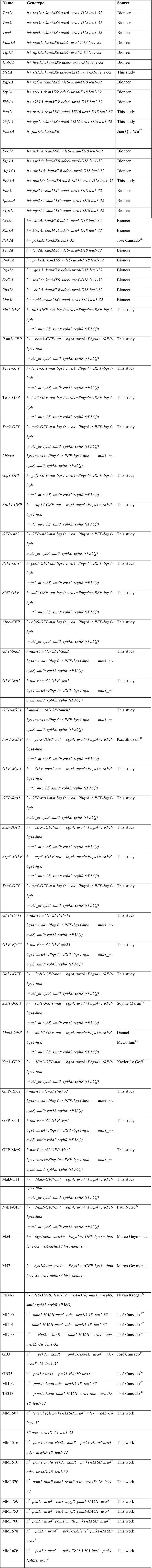
Cell lines used in this study.

**Supplementary table S2.**
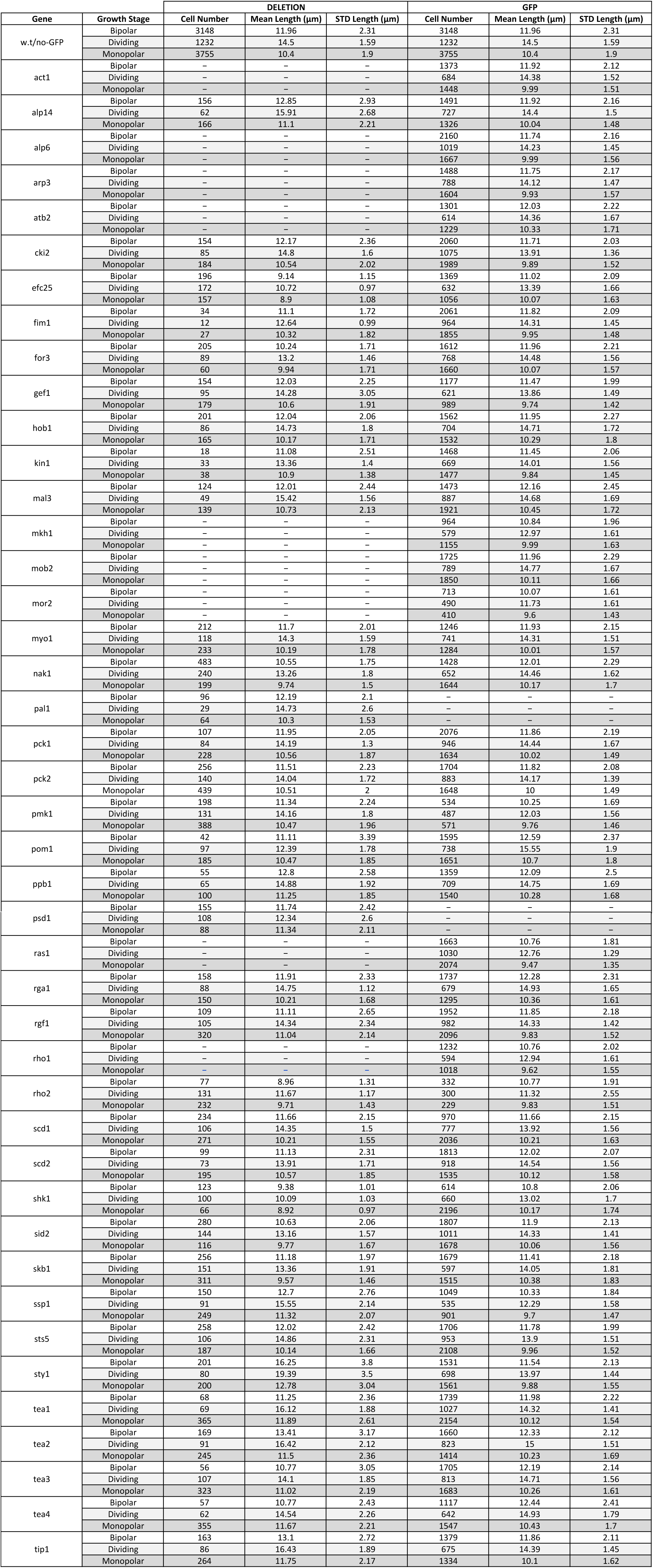

## REFERENCES

1. Barabási, A.-L. & Oltvai, Z. N. Network biology: understanding the cell’s functional organization. Nat. Rev. Genet. 5, 101–113 (2004).

2. Vidal, M., Cusick, M. E. & Barabasi, A.-L. Interactome networks and human disease. Cel. 144, 986–998 (2011).

3. Rolland, T. et al. A proteome-scale map of the human interactome network. Cell. 159, 1212–1226 (2014).

4. Costanzo, M. et al. The genetic landscape of a cell. Scienc. 327, 425–431 (2010).

5. Roguev, A. et al. Quantitative genetic-interaction mapping in mammalian cells. Nature method. 10, 432–437 (2013).

6. Laufer, C., Fischer, B., Billmann, M., Huber, W. & Boutros, M. Mapping genetic interactions in human cancer cells with RNAi and multiparametric phenotyping. Nature method. 10, 427–431 (2013).

7. Bassik, M. C. et al. A systematic mammalian genetic interaction map reveals pathways underlying ricin susceptibility. Cel. 152, 909–922 (2013).

8. Neumann, B. et al. Phenotypic profiling of the human genome by time-lapse microscopy reveals cell division genes. Natur. 464, 721–727 (2010).

9. Collinet, C. et al. Systems survey of endocytosis by multiparametric image analysis. Natur. 464, 243–249 (2010).

10. Hasson, S. A. et al. High-content genome-wide RNAi screens identify regulators of parkin upstream of mitophagy. Natur. 504, 291–295 (2013).

11. Graml, V. et al. A genomic multi-process survey of the machineries that control and link cell shape, microtubule organisation and cell cycle progression. Dev Cel.

12. La Carbona, S., Le Goff, C. & Le Goff, X. Fission yeast cytoskeletons and cell polarity factors: connecting at the cortex. Biol Cel. 98, 619–631 (2006).

13. Martin, S. G. Microtubule-dependent cell morphogenesis in the fission yeast. Trends Cell Bio. 19, 447–454 (2009).

14. Hayles, J. & Nurse, P. A journey into space. Nat Rev Mol Cell Bio. 2, 647–656 (2001).

15. Vaggi, F. et al. Linkers of cell polarity and cell cycle regulation in the fission yeast protein interaction network. PLoS Comput Bio. 8, e1002732 (2012).

16. Cortes, J. C. G. et al. The novel fission yeast (1,3)beta-D-glucan synthase catalytic subunit Bgs4p is essential during both cytokinesis and polarized growth. J Cell Sc. 118, 157–174 (2005).

17. Kim, D.-U. et al. Analysis of a genome-wide set of gene deletions in the fission yeast Schizosaccharomyces pombe. Nat. Biotechnol. 28, 617–623 (2010).

18. Benjamini, Y. & Hochberg, Y. Controlling the False Discovery Rate: A Practical and Powerful Approach to Multiple Testing on JSTOR. Journal of the Royal Statistical Society Series.… (1995).

19. Levin, D. E. & Bishop, J. M. A putative protein kinase gene (kin1+) is important for growth polarity in Schizosaccharomyces pombe. Proc Natl Acad Sci US. 87, 8272–8276 (1990).

20. Toya, M., Motegi, F., Nakano, K., Mabuchi, I. & Yamamoto, M. Identification and functional analysis of the gene for type I myosin in fission yeast. Genes Cell. 6, 187–199 (2001).

21. Mitchison, J. M. & Nurse, P. Growth in cell length in the fission yeast Schizosaccharomyces pombe. (1985).

22. Wixon, J. Featured organism: Schizosaccharomyces pombe, the fission yeast. Comp. Funct. Genomic. 3, 194–204 (2002).

23. Huisman, S. M. & Brunner, D. Cell polarity in fission yeast: a matter of confining, positioning, and switching growth zones. Semin. Cell Dev. Biol. 22, 799–805 (2011).

24. Wood, V. et al. PomBase: a comprehensive online resource for fission yeast. Nucleic Acids Re. 40, D695–9 (2012).

25. Moreno, S., Klar, A. & Nurse, P. Molecular genetic analysis of fission yeast Schizosaccharomyces pombe. Meth. Enzymol. 194, 795–823 (1991).

26. Perez, P. & Rincón, S. A. Rho GTPases: regulation of cell polarity and growth in yeasts. Biochem. J. 426, 243–253 (2010).

27. Madrid, M. et al. Stress-induced response, localization, and regulation of the Pmk1 cell integrity pathway in Schizosaccharomyces pombe. J Biol Che. 281, 2033–2043 (2006).

28. Soto, T. et al. Rga4 modulates the activity of the fission yeast cell integrity MAPK pathway by acting as a Rho2 GTPase-activating protein. J Biol Che. 285, 11516–11525 (2010).

29. Nunez, L. R. et al. Cell wall integrity MAPK pathway is essential for lipid homeostasis. J Biol Che. 283, 34204–34217 (2008).

30. Yu, J., Smith, V. A., Wang, P. P., Hartemink, A. J. & Jarvis, E. D. Advances to Bayesian network inference for generating causal networks from observational biological data. Bioinformatic. 20, 3594–3603 (2004).

31. Sachs, K., Perez, O., Pe’er, D., Lauffenburger, D. A. & Nolan, G. P. Causal protein-signaling networks derived from multiparameter singlecell data. Scienc. 308, 523–529 (2005).

32. Chatr-Aryamontri, A. et al. The BioGRID interaction database: 2015 update. Nucleic Acids Re. 43, D470–8 (2015).

33. Tong, A. H. Y. et al. Global mapping of the yeast genetic interaction network. Scienc. 303, 808–813 (2004).

34. TRANSWESD: inferring cellular networks with transitive reduction. 26, 2160–2168 (2010).

35. Zhou, F., Malher, S. & Toivonen, H. 2010 IEEE International Conference on Data Mining. 2010 IEEE 10th International Conference on Data Mining (ICDM. 659–668 (2010).doi:10.1109/ICDM.2010.133

36. Wachsmuth, M. et al. High-throughput fluorescence correlation spectroscopy enables analysis of proteome dynamics in living cells. Nat. Biotechnol. 33, 384–389 (2015).

37. Neumann, B. et al. High-throughput RNAi screening by time-lapse imaging of live human cells. Nature method. 3, 385–390 (2006).

38. Horn, T. et al. Mapping of signaling networks through synthetic genetic interaction analysis by RNAi. Nature method. 8, 341–346 (2011).

39. Bähler, J. & Pringle, J. R. Pom1p, a fission yeast protein kinase that provides positional information for both polarized growth and cytokinesis. Genes Dev. 12, 1356–1370 (1998).

40. Bähler, J. et al. Heterologous modules for efficient and versatile PCR-based gene targeting in Schizosaccharomyces pombe. Yeas. 14, 943–951 (1998).

41. Sato, M., Dhut, S. & Toda, T. New drug-resistant cassettes for gene disruption and epitope tagging in Schizosaccharomyces pombe. Yeas. 22, 583–591 (2005).

42. Sato, M., Toya, M. & Toda, T. Visualization of fluorescence-tagged proteins in fission yeast: the analysis of mitotic spindle dynamics using GFP-tubulin under the native promoter. Methods Mol. Biol. 545, 185–203 (2009).

43. Sato, M. & Toda, T. Alp7/TACC is a crucial target in Ran-GTPase-dependent spindle formation in fission yeast. Natur. 447, 334–337 (2007).

44. Riedl, J.. et al. Lifeact: a versatile marker to visualize F-actin. Nature method. 5, 605–607 (2008).

45. Arellano, M. et al. Schizosaccharomyces pombe protein kinase C homologues, pck1p and pck2p, are targets of rho1p and rho2p and differentially regulate cell integrity. J Cell Sc. 112(Pt 20), 3569–3578 (1999).

46. Schneider, C. A., Rasband, W. S. & Eliceiri, K. W. NIH Image to ImageJ: 25 years of image analysis. Nature method. 9, 671–675 (2012).

47. Wu, J. Q., Bahler, J. & Pringle, J. R. Roles of a fimbrin and an alpha-actinin-like protein in fission yeast cell polarization and cytokinesis. Mol Biol Cel. 12, 1061–1077 (2001).

48. Tatebe, H., Nakano, K., Maximo, R. & Shiozaki, K. Pom1 DYRK regulates localization of the Rga4 GAP to ensure bipolar activation of Cdc42 in fission yeast. Curr Bio. 18, 322–330 (2008).

49. Bendezú, F. O. & Martin, S. G. Cdc42 explores the cell periphery for mate selection in fission yeast. Curr Bio. 23, 42–47 (2013).

50. Hou, M.-C., Wiley, D. J., Verde, F. & McCollum, D. Mob2p interacts with the protein kinase Orb6p to promote coordination of cell polarity with cell cycle progression. J Cell Sc. 116, 125–135 (2003).

51. Cadou, A. et al. Kin1 is a plasma membrane-associated kinase that regulates the cell surface in fission yeast. Mol Microbio. 77, 1186–1202 (2010).

52. Leonhard, K. & Nurse, P. Ste20/GCK kinase Nak1/Orb3 polarizes the actin cytoskeleton in fission yeast during the cell cycle. J Cell Sc. 118, 1033–1044 (2005).

53. Roguev, A., Wiren, M., Weissman, J. S. & Krogan, N. J. High-throughput genetic interaction mapping in the fission yeast Schizosaccharomyces pombe. Nature method. 4, 861–866 (2007).

54. Barba, G. et al. Activation of the cell integrity pathway is channelled through diverse signalling elements in fission yeast. Cell. Signal. 20, 748–757 (2008).

